# Viola: a structural variant signature extractor with user-defined classifications

**DOI:** 10.1101/2021.03.31.437648

**Authors:** Itsuki Sugita, Shohei Matsuyama, Hiroki Dobashi, Daisuke Komura, Shumpei Ishikawa

## Abstract

Here, we present Viola, a Python package that provides structural variant (SV; large scale genome DNA variations that can result in disease, e.g., cancer) signature analytical functions and utilities for custom SV classification, merging multi-SV-caller output files, and SV annotation. We demonstrate that Viola can extract biologically meaningful SV signatures from publicly available SV data for cancer and we evaluate the computational time necessary for annotation of the data.

**Availability:** Viola is available on pip (https://pypi.org/project/Viola-SV/) and on GitHub (https://github.com/dermasugita/Viola-SV).

## 1. Introduction

Somatic mutations in cancer are the cumulative result of DNA aberrations caused by diverse mutational processes. Recently, large scale studies of human cancer have revealed characteristic patterns of mutation types, i.e., mutational signatures, arising from specific processes of single nucleotide variant formation. These studies often provide theoretical explanations for known mutational processes and their consequences, e.g., C>A substitutions and CC>TT alterations caused by smoking and ultraviolet light exposure, respectively.

Structural variants (SVs) are another type of DNA mutation, defined as events larger than 50-bp in size or involving multiple chromosomes, occupying non-negligible proportions of mutations in cancer cells (Mills *et al.*, 2011; Yi and Ju, 2018). Signature analysis of SVs may potentially provide novel insights into carcinogenesis. The development of high-throughput sequencing technologies and powerful SV callers has improved the accuracy of SV event identification. Several mechanisms of SV formation have also been identified (Yi and Ju, 2018). Therefore, research on SV signatures is gradually becoming realistic.

To date, several attempts have been made to decompose SV patterns into SV signatures, but an established method has yet to be realized. Previous studies have mainly classified SVs according to segment size and revealed an association between small tandem duplications and BRCA1 mutations (Li *et al.*, 2020; Nik-Zainal *et al.*, 2016). However, a consensus has not been achieved on a precise SV classification method.

SVs can be classified by metrics other than length. Li *et al.* (2020) also used replication timing and common fragile sites (CFSs). Interestingly, the biological meaningfulness of replication timing and CFSs has been reported, e.g., the signatures of medium-sized (50–500 kb) tandem duplications occurring at the site of late replication timing have been associated with CDK12 driver mutations, whereas CFS signatures have been associated with gastrointestinal cancer. Other SV classification methods, such as microhomology and association of transposons, have yet to be considered in detail; therefore, further analysis is required to identify a suitable SV classification method for signature analysis.

At present, very few tools are available for SV signature analysis. To the best of our knowledge, pyCancerSig (Thutkawkorapin *et al.*, 2020), which is the first tool that can handle SVs for cancer mutation signature analysis, is the only SV signature analysis tool currently available. However, pyCancerSig has limitations in SV classifications as it only supports traditional SV classes, i.e., deletion, duplication, inversion, and translocation, and length-based classification.

The time-consuming nature of parsing variant call format (VCF) files is also an obstacle to SV analysis. VCF is the de facto standard format by which genetic variant data are recorded with high human readability. However, from a data management perspective, VCF can be a bottleneck for analysis owing to its complex structure. For SVs in particular, accurate interpretation of VCF records at the single nucleotide level requires considerable learning costs. Difficulties with VCF interpretation cannot be ignored because even a 1-bp error in positioning SVs can have critical consequences, e.g., in microhomology analysis.

Merging SV calls from different callers is also an issue in SV analysis. Precision of SV detection can be improved by merging the results of multiple SV callers (Cameron *et al*, 2019; Kuzniar *et al.*, 2020); however, different SV callers use different ways to represent VCF files, which makes integration challenging.

Here, we present Viola, a highly customizable and flexible Python package that supports SV signature analysis with user-defined SV classification, matrix-generation functions, and a file exportation system that is compatible with external statistical utilities and facilitates interpretation of results. Viola accepts VCF files from four popular SV callers, namely Manta, Delly, Lumpy, and Gridss, and can also read BEDPE format (Cameron. *et al.*, 2017; Chen *et al.*, 2016; Layer *et al.*, 2014; Rausch *et al.*, 2012). Viola also provides an intuitive VCF file manager for filtering, annotating, converting VCF to BEDPE, and multicaller merging.

## 2. Implementation

### 2.1 Data Structure

Viola converts input SV data files, such as VCF and BEDPE files, into our original Python classes. Instances of these classes store SV data as a set of tidy rectangular tables linked via identifiers such as SV ID output by the SV callers (Supplementary Figure S1). These tables follow the principles of tidy data, i.e., each SV record is a row, each variable is a column, and each type of observational unit is a table (Wickham, 2014). Consequently, storage of multiple values in one element is avoided, in contrast to the INFO and FORMAT columns of a VCF file. Hence, a specific single value can be accessed by simply specifying the row and column of the table of interest; this provides freedom in data handling without the need for cumbersome codes.

### 2.2 User Interface

Viola is written in the Python. Although it is intended for use within Python scripts, some features are available from the command line.

Viola supports SV signature analysis with user-defined SV classes (Figure 1A; Supplementary Figure S1A, B). A simple feature matrix based on traditional SV types and SV length, output by the SV caller can be generated from the command line. Advanced uses such as annotation, filtering, and multicaller intersection, which are required to generate a complex feature matrix, are supported within Python scripts. In combination with these functions, it is possible to define a wide variety of SV classes, such as “Duplications located on CFS sites” and “Deletions less than 50 kb in size, located on the early replication timing zones.” These operations can be implemented with simple syntax and are designed to refine the SV classification by trial and error (Supplementary Figure S2B).

**Fig. 1.**
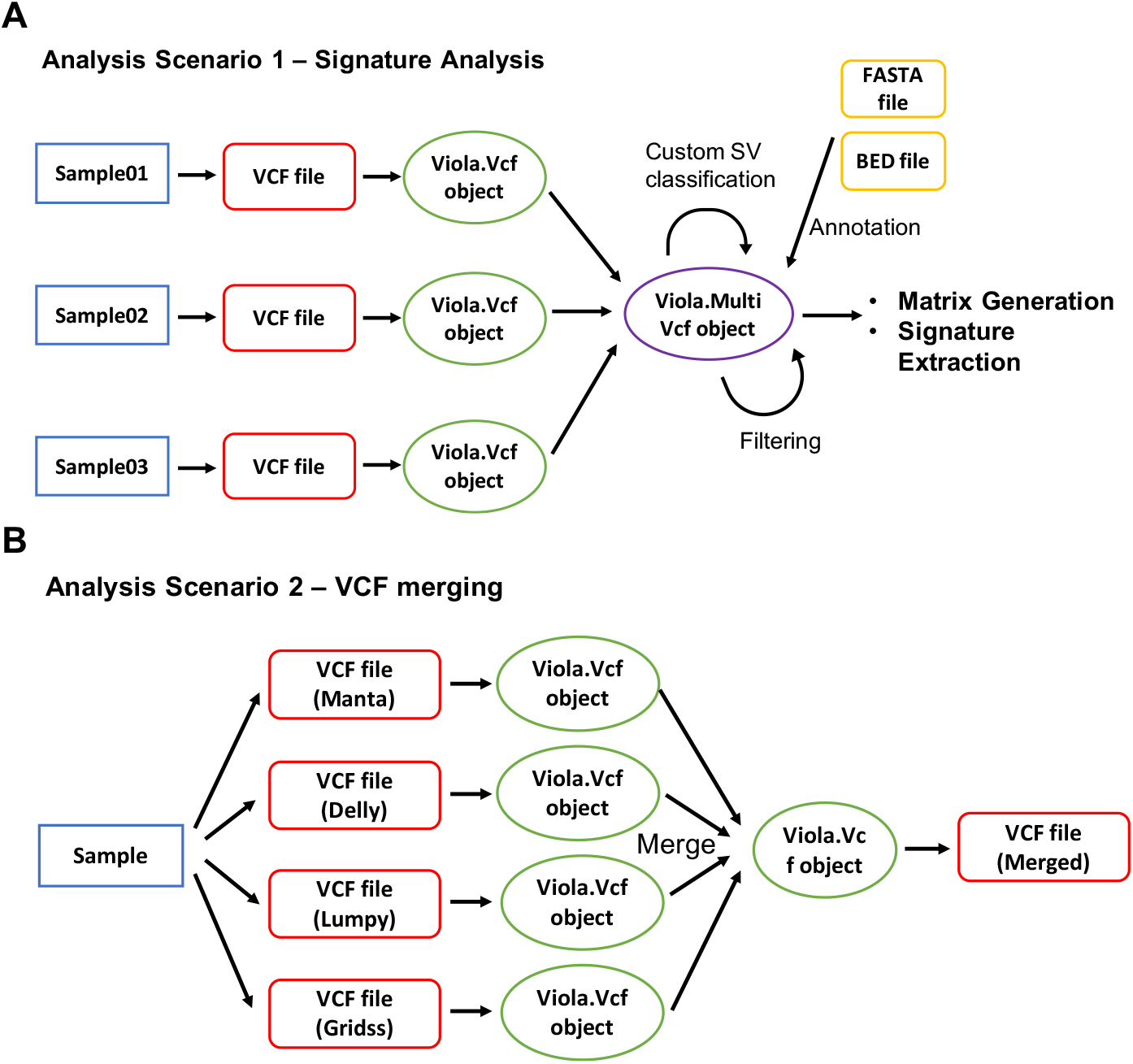
Visualization of the data flow in the main analysis scenarios. **(A)** Process of feature matrix generation from multiple samples. **(B)** Overview of VCF merging system.

From an internal data structure perspective, user-defined SV classes are interpreted as new INFO entries of the VCF file. Hence, users can output new VCF or BEDPE files with annotation of novel SV classes as well as generating a signature-analysis-ready feature matrix according to these additional SV classes.

Alongside signature analysis, Viola has the following features:

- Support of well-known SV callers including Manta, Delly, Lumpy, and Gridss. The notation has been unified as much as possible to facilitate subsequent processing including merging (Figure 1B).
- Fast annotation methods that utilize the interval tree algorithm. Source files in BED format are acceptable; thus, information such as gene names, CFSs, replication timing, and copy number can be annotated if they can be expressed in BED format.
- An intuitive method for filtering SV records. In addition to filtering for genomic coordinates and INFO fields, filtering for FORMAT fields is possible.
- Estimations of the length and sequence of microhomology from SV breakpoint positions. Where SV callers do not return microhomology information or publicly available SV data does not contain such information, Viola can estimate microhomology using the reference sequence.

The use of these characteristics is described in detail in the official Viola documentation, which is available online (https://dermasugita.github.io/ViolaDocs/docs/html/index.html).

### 2.3 Custom SV Classification Overview

With Viola, any information in the INFO field of the VCF can be used for SV classification. Many SV callers write the SV type and length in the INFO field by default making it easy to classify by these variables. For BEDPE files that do not define a field corresponding to the INFO field in a VCF file, Viola will automatically generate INFO fields such as SV length and type. Additionally, new INFO fields can be added using BED file annotation and microhomology prediction. BED files can be used to annotate genes, CFSs, replication timing, copy numbers, etc., which individually or in combination can be used to classify SVs.

## 3. Application

### 3.1 Matrix Generation with Simple Code

We ran Viola to generate an SV feature matrix using public BEDPE files reported in a PCAWG study (Li *et al.*, 2020). First, we downloaded 2,748 BEDPE files from the ICGC data portal and used Viola to read 2,605 of these files that were not empty as a MultiBedpe instance. Second, the instance was successfully annotated by CFSs and replication timing BED files that we built according to the PCAWG study. We defined 25 SV classes according to CFSs, replication timing, and SV length and then generated a 2,605 × 25 feature matrix. These operations were written in only 11 lines of the Python code, excluding code for custom SV definitions (Supplementary Figure S2A). The matrix generated here can be easily reproduced by following the tutorial in the Viola official document.

### 3.2 Signature Extraction Analysis

We extracted nine SV signatures from the generated matrix using a function of Viola that simultaneously performs non-negative matrix factorization (NMF) and cluster stability evaluation (Supplementary Figures S3 and S4). Several signatures, including the signatures of CFSs, small deletions (<50 kb), and small duplications (<50 kb), were comparable to those in the PCAWG study (Li *et al.*, 2020). We further explored the association between each of the nine signatures and driver mutations of three well-known DNA repair genes: *BRCA1, BRCA2*, and *CDK12* (Supplementary Table S1). These genes were significantly associated with the small duplication signature, small deletion signature, and medium-large duplication signature, as expected from previous studies (Li *et al.*, 2020; Menghi *et al.*, 2018; Nik-Zainal *et al.*, 2016; Popova *et al.*, 2016) (Supplementary Table S1).

### 3.3 Multicaller VCF Merging

We synthesized VCF files that mimicked the output from Manta, Delly, Lumpy, and Gridss. These files shared several SVs recorded with errors within 100 bp of each other. Four VCF files were read as the object of Viola and then merged, with 100 bp being specified as the option for proximity. The identifier was added as a new INFO and the same SVs were given the same ID. We removed SV records called by only one SV caller. Finally, all shared SVs were merged as expected and successfully exported as a VCF file (Supplementary Data 1).

### 3.4 Annotation Performance

We tested the performance of the annotations on 2,605 BEDPE files using 18 lines of CFS BED files. In total, 618,492 break-ends were annotated according to whether each was present or absent on the CFS. On average, this took 7.5 min to complete using a single thread on an Ubuntu x86_64 server (Intel Core i7-8700K CPU at 3.70 GHz).

## 4. Conclusion

We developed Viola, a tool for SV signature analysis that allows highly customizable SV classification. This tool also overcomes the difficulty of parsing current VCF files as well as the problem of different notations derived from different callers. Viola will help stimulate research in the SV field to better understand the biological significance of SVs.

## Acknowledgements

We thank Enago (www.enago.jp) for the English language review.

## Supplementary Information

**Supplementary Figure S1.**
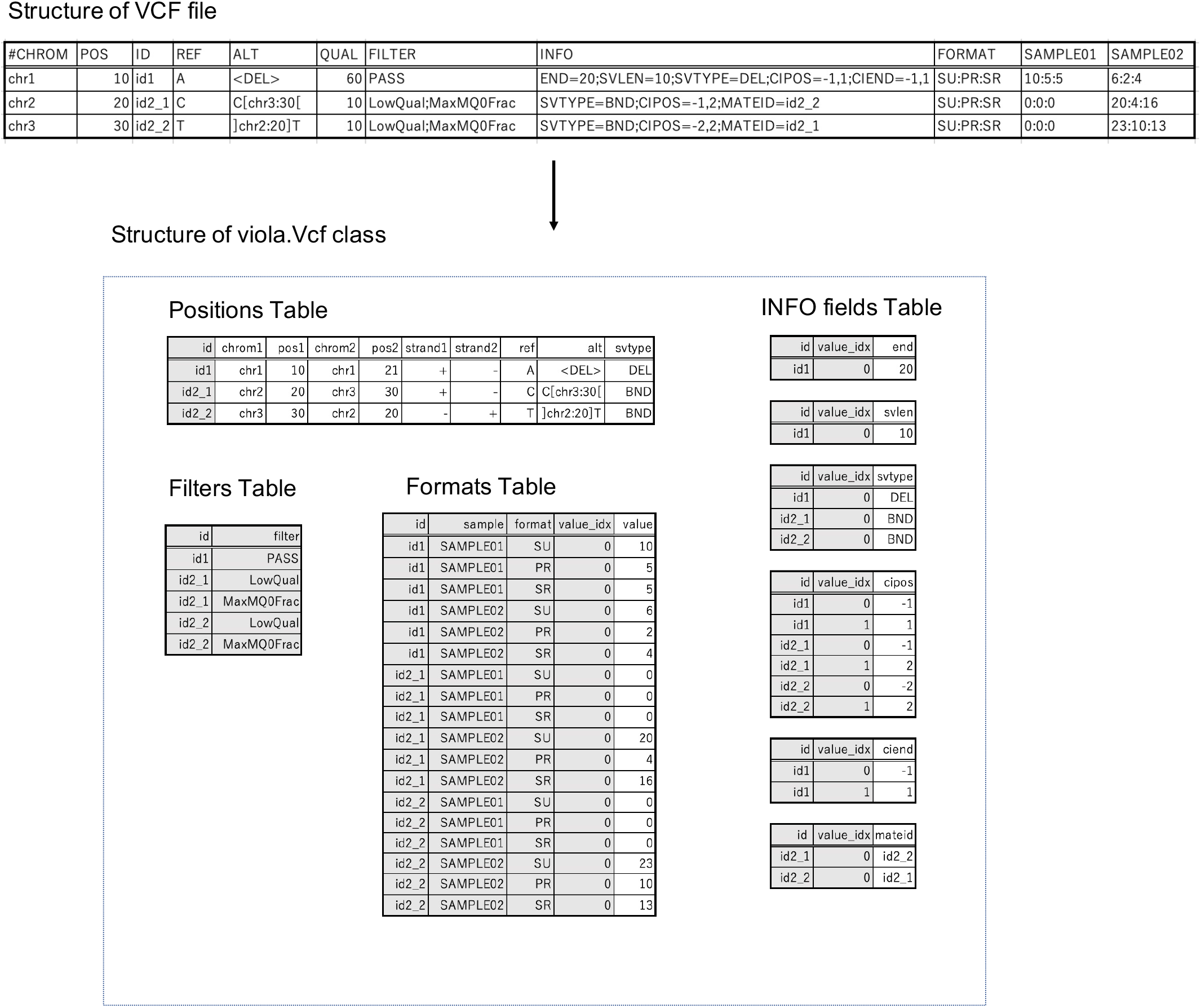
Data structure of a viola.Vcf object. The upper part of the figure shows an example of a Manta-like VCF. As shown in the lower part of the figure, the viola.Vcf object holds the information of a VCF file in several rectangular tables. The tables are related to each other by VCF IDs. The grey columns are the primary key or composite primary key of the table. The header information of the VCF is also stored as tables (not shown). Abbreviations: POS: start position of the SV; END: end position of the SV; SVLEN: length of the SV; SVTYPE: type of SV; CIPOS: confidence interval around POS; CIEND: confidence interval around END; MATEID: ID of mate break end; SU: count of supporting reads of the SV; PR: count of paired end reads supporting the SV; SR: count of split reads supporting the SV.

**Supplementary Figure S2.**
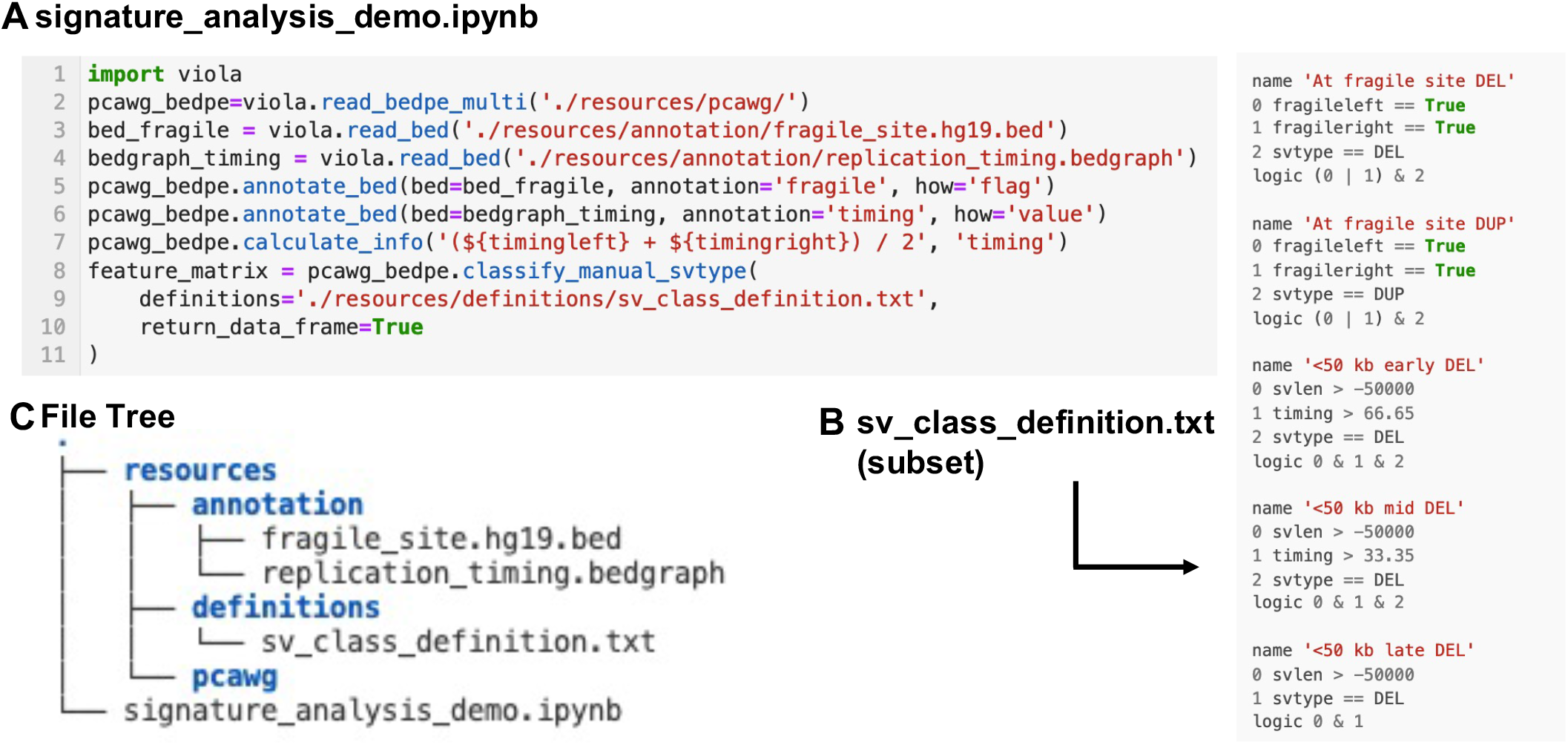
Example code for feature matrix generation. **(A)** (1) Import Viola package. (2) Read BEDPE files under the “pcawg” directory as viola.MultiBedpe object. (3 and 4) Load common fragile site and replication timing BED/BEDGRAPH for annotation*. (5 and 6) Annotate “pcawg_bedpe” variable with the BED/BEDGRAPH loaded above. (7) Obtain mean replication timing for each SV breakpoint. (8–11) Classify custom SV type according to the definition file and export feature matrix. **(B)** Definition file for custom SV classification. Each SV class definition consists of a line specifying the SV class name, lines describing the conditions, and a line passing the set operation of the conditions. Note that the file content shown here is part of all SV definitions used in this study. **(C)** File tree of this analysis. * Currently, a clear distinction between BED and BEDGRAPH files is not made in relation to the annotation of Viola objects since only the first four columns of these files are used for annotation purposes.

### Signature Extraction Procedure

Here, we describe how SV signatures were extracted from the PCAWG dataset. To determine the number of signatures, *K*, we evaluated the stability of signatures derived from non-negative matrix factorization (NMF) and its reconstruction error. Detailed steps are provided below.

1. Generate a new 2,605 × 25 matrix, 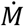, by bootstrapping the original matrix M. Here each element 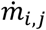 of 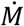 is chosen with a probability of *m_i,j_*/Σ_*i,j*_*m_i,j_*, where *m_i,j_* is each element of M 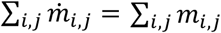.
2. Apply NMF to the bootstrapped matrix 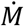 to obtain an exposure matrix, 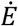 with 2,605 × *K* and a signature matrix, 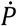, with *K* × 25. 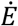 and 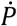 are initialized by a non-negative double singular decomposition method with zeros filled with the average of 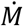. Kullback–Leibler divergence is used for loss function.
3. Perform step 1 and 2 for 100 iterations to obtain 100K signatures.
4. Use a K-means method for clustering 100K signatures into K clusters with the constraint that signatures from the same iteration should not been assigned to the same cluster. The average silhouette score is calculated for stability evaluation.
5. The average signature matrix 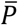 is constructed with *K* × 25. Each row of 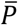 is the centroid of the K-means clustering performed in step 4. The average exposure matrix 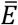 is then calculated by NMF using the original matrix M and 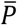, where the matrix 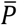 is not updated while NMF. Finally, the Kullback–Leibler divergence of M and 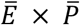 was calculated as reconstruction error.

Steps 1–5 were conducted for each *K* ranging from 2 to 13 (Supplementary Figure S3).

**Supplementary Figure S3.**
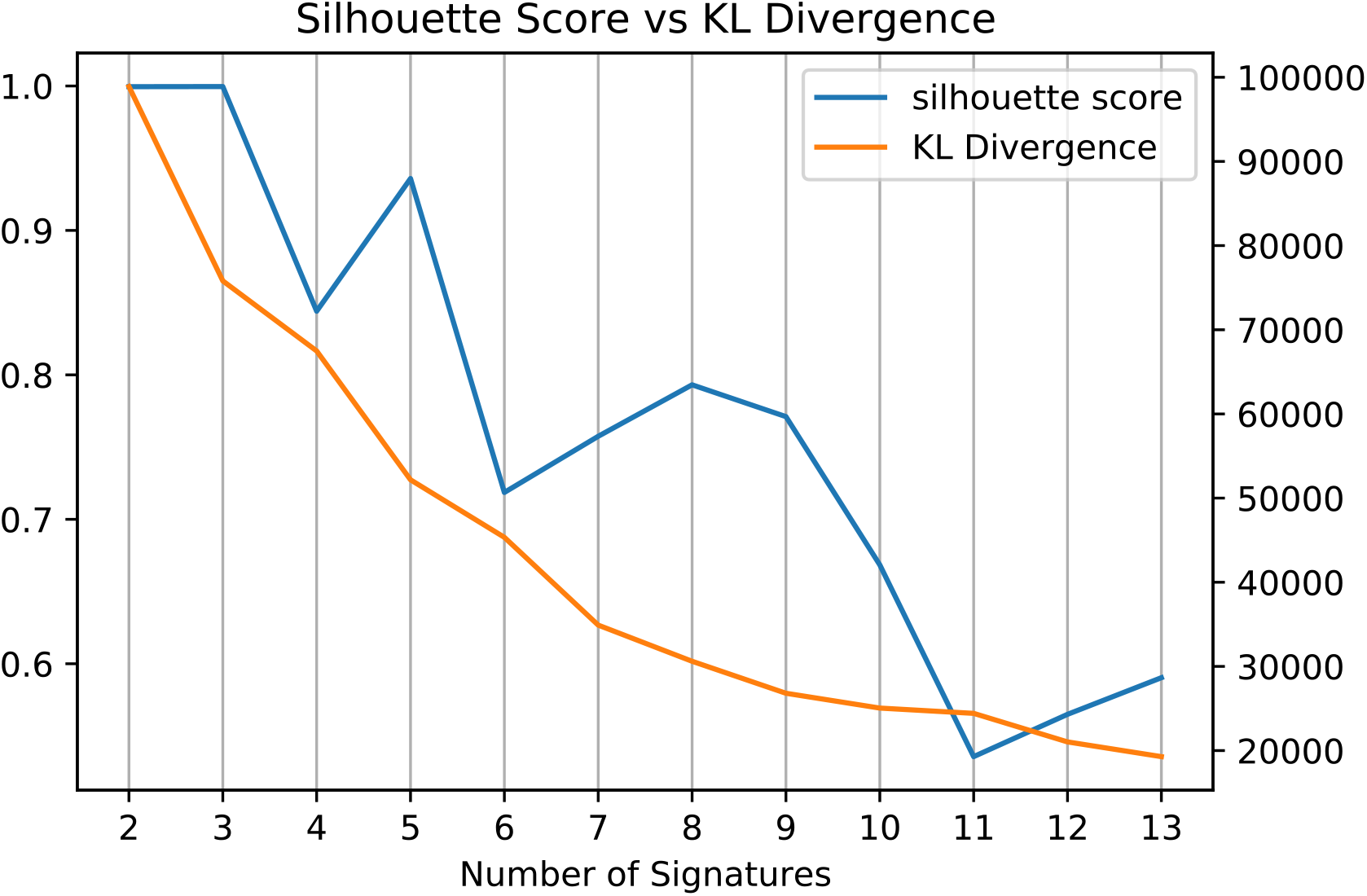
Average silhouette score of K-means clusters and reconstruction error for the number of signatures (K). After a manual assessment of each K with reference to the stability score and reconstruction error, we chose K = 9 as the number of signatures. Extracted signatures are shown in Supplementary Figure S4.

**Supplementary Figure S4.**
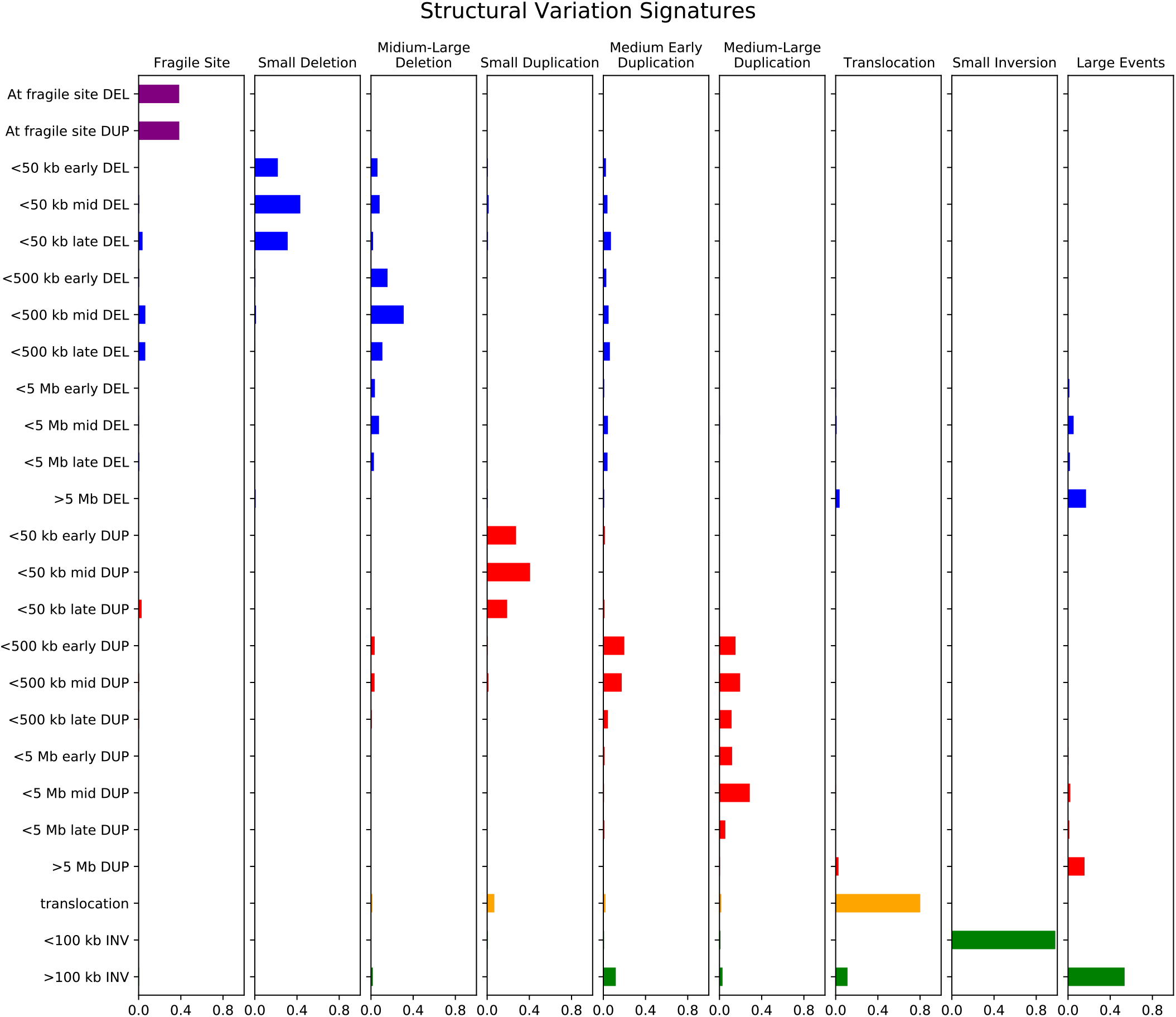
Nine signatures extracted from the PCAWG dataset using Viola.

### Statistical testing of the association between signatures and driver mutations

We obtained several signatures that were comparable with those in the PCAWG report such as the small deletion signature and medium-large duplication signature. Statistical significance was tested for the effect of driver mutations in BRCA1, BRCA2, and CDK12 on the nine signatures. The p value of each signature was calculated using a linear model that considered the histological type of each sample (Supplementary Table S1).

**Supplementary Table S1.** Statistical significance of the effect of driver mutations in *BRCA1, BRCA2*, and *CDK12* on nine signatures. Negative log *p* values are shown (\**p* < 0.01).

|  | *BRCA1* | *BRCA2* | *CDK12* |
| --- | --- | --- | --- |
| **Fragile Site** | 0.205 | 0.527 | 0.192 |
| **Small Deletion** | 0.084 | **23.278\*** | 0.750 |
| **Medium-Large Deletion** | 0.401 | 1.142 | 0.036 |
| **Small Duplication** | **26.030\*** | 1.579 | 1.055 |
| **Medium Early Duplication** | 0.389 | 0.421 | 0.057 |
| **Medium-Large Duplication** | 0.663 | 1.526 | **6.251\*** |
| **Translocation** | 0.122 | 0.218 | 0.950 |
| **Small Inversion** | 0.729 | 0.520 | 0.128 |
| **Large Events** | **2.877\*** | **2.042\*** | 0.479 |

